# Optimized tRNA structure–seq reveals robust tRNA secondary structures in *S. cerevisiae* under mild stress conditions

**DOI:** 10.64898/2026.02.24.707850

**Authors:** Komei Yanagihara, Futa Konishi, Teppei Matsuda, Akira Hirata, Hiroyuki Hori, Philip C. Bevilacqua, Ryota Yamagami

**Affiliations:** Department of Applied Chemistry, Graduate School of Science and Engineering, Ehime University, 3 Bunkyo-cho, Matsuyama, Ehime 790-8577, Japan; Department of Natural Science, Division of Science and Technology, Graduate School of Sciences and Technology for Innovation, Tokushima University, 2-1 Minamijosanjima-cho, Tokushima, Tokushima 770-8506, Japan; Department of Chemistry, Pennsylvania State University, University Park, PA 16802, USA; Center for RNA Molecular Biology, Pennsylvania State University, University Park, PA 16802, USA; Department of Biochemistry and Molecular Biology, Pennsylvania State University, University Park, PA 16802, USA

**Author notes:** **Corresponding author:** Ryota Yamagami, Address: 3 Bunkyo-cho, Matsuyama, Ehime 790-8577, Japan.

**Keywords:** tRNA, tRNA modification, tRNA structure, prediction of RNA structure, mutation profiling, stress

## Abstract

RNA structure plays a crucial role in diverse biological processes beyond the translation of genetic information. Therefore, the development of reliable methods for RNA structure prediction is essential for understanding RNA structure–related functions, however accurate and comprehensive RNA structure prediction remains challenging. Here, we focus on secondary structure prediction of transfer RNA (tRNA) using structure probing coupled with next–generation sequencing (tRNA Structure–seq). In silico prediction of *Saccharomyces cerevisiae* tRNA secondary structures achieves only 56.9% accuracy on average. Incorporation of dimethyl sulfate (DMS) probing data improve prediction accuracy to 87.4%, which is still not sufficient for practical tRNA structure prediction. To overcome this, we optimized the tRNA Structure–seq analysis pipeline by explicitly incorporating natural tRNA modifications detected in tRNA sequencing data and by refining pseudo–free energy parameters specifically optimized for tRNA structure prediction. Using this optimized pipeline, the average prediction accuracy is remarkably improved to 94%. Furthermore, analysis of multiple structural conformations predicted from DMS probing data indicates that *S. cerevisiae* tRNAs predominantly adopt the canonical cloverleaf secondary structure under *in vivo* conditions. Finally, we examined tRNA structures under mild stress conditions, including heat stress, osmotic stress, and antibiotic stress. These perturbations had minimal effects on in vivo tRNA secondary structure, demonstrating that *S. cerevisiae* tRNAs maintain structural stability under physiologically relevant stress conditions. In summary, our results establish an optimized tRNA Structure–seq analysis that enables highly accurate tRNA secondary structure prediction and reveals the intrinsic robustness of tRNA structures in living cells.

## Introduction

In the classical view of the central dogma of genetic information, RNA was regarded primarily as a carrier of genetic information. In recent years, however, a wide variety of RNA functions beyond the translation of genetic information have been identified across the three domains of life, as well as in viruses. Many of these RNA functions are closely linked to RNA structure (Spitale and Incarnato 2023). For example, riboswitches bind cognate biological small molecules such as S–adenosyl–L–methionine, amino acids, metabolites, or nucleic acids. Ligand binding induces dynamic structural rearrangements in riboswitches, enabling the RNA to regulate transcription, translation, or splicing (Kavita and Breaker 2023). RNA thermometers provide another representative example of structure–dependent regulation. These RNAs act as a thermosensor by altering their secondary structures in response to temperature changes, thereby exposing or sequestering key regulatory elements such as ribosome–binding sites (Kortmann and Narberhaus 2012; Jolley et al. 2023). Also, transfer RNAs (tRNAs) have long been recognized to be critical to translating the genetic code but more recently have been shown to regulate gene expression (Su et al. 2020), refold during stress (Yamagami et al. 2022), and be effective in RNA medicine (Coller and Ignatova 2024). In this way, structure of diverse RNAs plays critical roles in diverse biological processes. Consequently, accurate prediction of RNA secondary structure is necessary for understanding these RNA structure–related regulatory functions.

Over the past four decades, numerous computational approaches have been developed to predict RNA secondary structures, most of which rely on identifying minimum free energy structures calculated with empirical thermodynamic parameters (Freier et al. 1986; Jaeger et al. 1989; Xia et al. 1998; Mathews et al. 1999; Christiansen and Znosko 2008; Turner and Mathews 2010). While the effort enables us to rapidly predict RNA secondary structure genome-wide, accurate prediction of complete RNA secondary structures in silico remains challenging.

To address this, RNA structure–sensitive chemical probing reagents such as dimethyl sulfate (DMS) and SHAPE reagents have been widely utilized for experimentally predicting RNA secondary structure (Moazed et al. 1986; Mathews et al. 2004; Spitale et al. 2013; Ding et al. 2014). DMS selectively modifies the N1 position of adenine, the N3 position of cytosine, and the N7 position of guanine, which results in the formation of 1–methyladenosine (m^1^A), 3–methylcytidine (m^3^C), and 7–methylguanosine (m^7^G), respectively. SHAPE reacts at 2’–hydroxyl group of the ribose backbone, reviewed in (Mitchell et al. 2019a). These modified nucleosides leave signals during reverse transcription (RT) that are typically detected as either RT stops or misincorporation at those positions through mutational profiling (MaP) (Siegfried et al. 2014). The RT stop signatures were traditionally detected by gel–based sequencing for a single RNA but now are read by next generation sequencing technologies for MaP for many RNAs. Importantly, Watson–Crick base pairing and stable secondary structure elements, as well as tertiary structure and binding of proteins, protect these reactive sites from chemical modification. Thus, DMS and SHAPE reactivities inversely reflect local base pairing status within an RNA helix (i.e. base pairing inhibits these chemical reactions). These RNA structural data from DMS and SHAPE probing experiments have improved prediction accuracy by integrating with computational RNA secondary structure prediction (Mathews et al. 2004; Merino et al. 2005; Deigan et al. 2009). Furthermore, the emergence of next–generation sequencing technologies have enabled transcriptome–wide detection of DMS and SHAPE reactivities from each transcript, which accelerated high–throughput prediction of in vivo RNA structures (Lucks et al. 2011; Ding et al. 2014; Rouskin et al. 2014; Siegfried et al. 2014; Spitale et al. 2015; Zubradt et al. 2017; Guo et al. 2020; Ritchey et al. 2020). Based on these technological advances, diverse analytical frameworks have been developed. For example, Rsample uses chemical reactivity data to estimate RNA secondary structure folding ensembles (Spasic et al. 2018), and the DREEM algorithm performs chemical reactivity–based clustering of high–throughput reads to infer multiple RNA conformations (Tomezsko et al. 2020). Single molecule–based Structure–seq approaches, which retain individual reactivity information in a single read rather than averaging reactivities across transcripts, can analyze multiple RNA conformations and their populations at a single–molecule level (Yang et al. 2022). In addition, chemicals that probe tertiary contacts have been used for analyzing three–dimensional RNA–RNA interactions (Christy et al. 2021).

In 2022, we developed a Structure–seq method specialized for the prediction of secondary structure of tRNA, which we call tRNA Structure–seq (Yamagami et al. 2022; Meyer et al. 2023a; Meyer et al. 2023b). Transfer RNA is one of the best studied RNA molecules, and its structure and functions have been extensively characterized. Transfer RNAs are composed of four stems: acceptor stem, D–stem, anticodon stem, and T–stem, and fold into a canonical cloverleaf secondary structure (Sprinzl et al. 1998) and an L–shaped tertiary structure (Shi and Moore 2000). Transfer RNA undergoes diverse post–transcriptional modifications that regulate tRNA structure, function, and decoding system during protein synthesis (Suzuki 2021). Our method utilizes DMS–modified tRNA and a highly processive reverse transcriptase originally isolated from a bacterial group II intron, for reverse transcription with detection by MaP (Zhao et al. 2018; Guo et al. 2020). This reverse transcriptase introduces mutations at both DMS–modified and naturally modified nucleoside positions. Incorporation of DMS reactivity data improved prediction accuracy of *Escherichia coli* tRNA secondary structures under in vivo conditions to approximately 95%. In addition, tRNA Structure–seq analysis under heat–stress conditions suggested that base pairs in the acceptor stem undergo dynamic breathing, and heat stress causes partial disruption of the acceptor stem in a subset of tRNAs (Yamagami et al. 2022). Although tRNA Structure–seq enables highly accurate prediction of tRNA secondary structure, the method has thus far been validated only in *E. coli*. In this study, we extend tRNA Structure–seq to the eukaryotic model organism *Saccharomyces cerevisiae* (*S. cerevisiae*) and further optimize the analysis to improve structure prediction accuracy and robustness of the method.

## Results

### tRNA Structure–seq successfully detected DMS modifications and natural modifications in *S. cerevisiae* tRNAs

To investigate the tRNA structurome in eukaryotes, we applied our tRNA Structure–seq technique to *S. cerevisiae*. The tRNA Structure–seq analysis is composed of six major steps: (Step 1) DMS treatment and tRNA isolation; (Step 2 – Step 4) library preparation and next–generation sequencing; (Step 5 and Step 6) calculation of DMS reactivity at a single–nucleotide resolution and prediction of secondary structure by RNAstructure Fold (Fig. 1A). During both in vitro and in vivo structure probing experiments for native *S. cerevisiae* tRNAs, plus/minus DMS conditions were performed. ShapeMapper2 processes these two datasets and identifies nucleotide positions with high mutation rates in the minus DMS data as possible natural modification sites. Using this strategy, tRNA Structure–seq successfully detected both DMS–induced and naturally modified nucleosides in *E. coli* tRNAs using Marathon reverse transcriptase in our previous study (Yamagami et al. 2022). In addition, our previous work demonstrated that DMS–modified guanosines can be detected when we used Mn²⁺ rather than Mg²⁺ during reverse transcription (Yamagami et al. 2022). In this study, we tested Induro reverse transcriptase, which is also derived from a bacterial group II intron and has recently been used in tRNA sequencing studies because of its high processivity on highly structured and heavily modified tRNAs and its ability to induce misincorporations at native modification sites (Nakano et al. 2025). Using Induro reverse transcriptase, we obtained highly reproducible DMS reactivity profiles across the three biological replicates analyzed in this study (Supplemental Figs. 1 and 2). In our previous work, we observed distinct mutation specificities at DMS–induced modification sites in *E. coli*. We reanalyzed that data and found that, at m^1^A sites, Marathon reverse transcriptase produced 59.3% A to T, 28.2% A to G, 10.4% A to C substitutions, and 2.1% indels (Supplemental Fig. 3A). We therefore wondered whether mutation specificities at DMS–modified nucleosides are conserved between Marathon and Induro reverse transcriptases. Comparison of the two enzymes revealed that mutation specificities at DMS–modified adenosines were highly similar between Marathon and Induro reverse transcriptases; compare lefthand pie charts in Supplemental Figures 3a and 3b. In contrast, mutation specificities at DMS–modified cytidines differed slightly between the two enzymes. Specifically, Marathon reverse transcriptase predominantly produced C to A (50.1%) and C to T (31.1%) substitutions, whereas Induro reverse transcriptase favored C to T (53.3%) followed by C to A (29.0%) substitutions under the in vivo condition (Supplemental Fig. 3A–3C). These results indicate that mutation specificity at DMS–modified nucleotides is somewhat influenced by the enzymatic properties of the reverse transcriptase.

**Figure 1.**
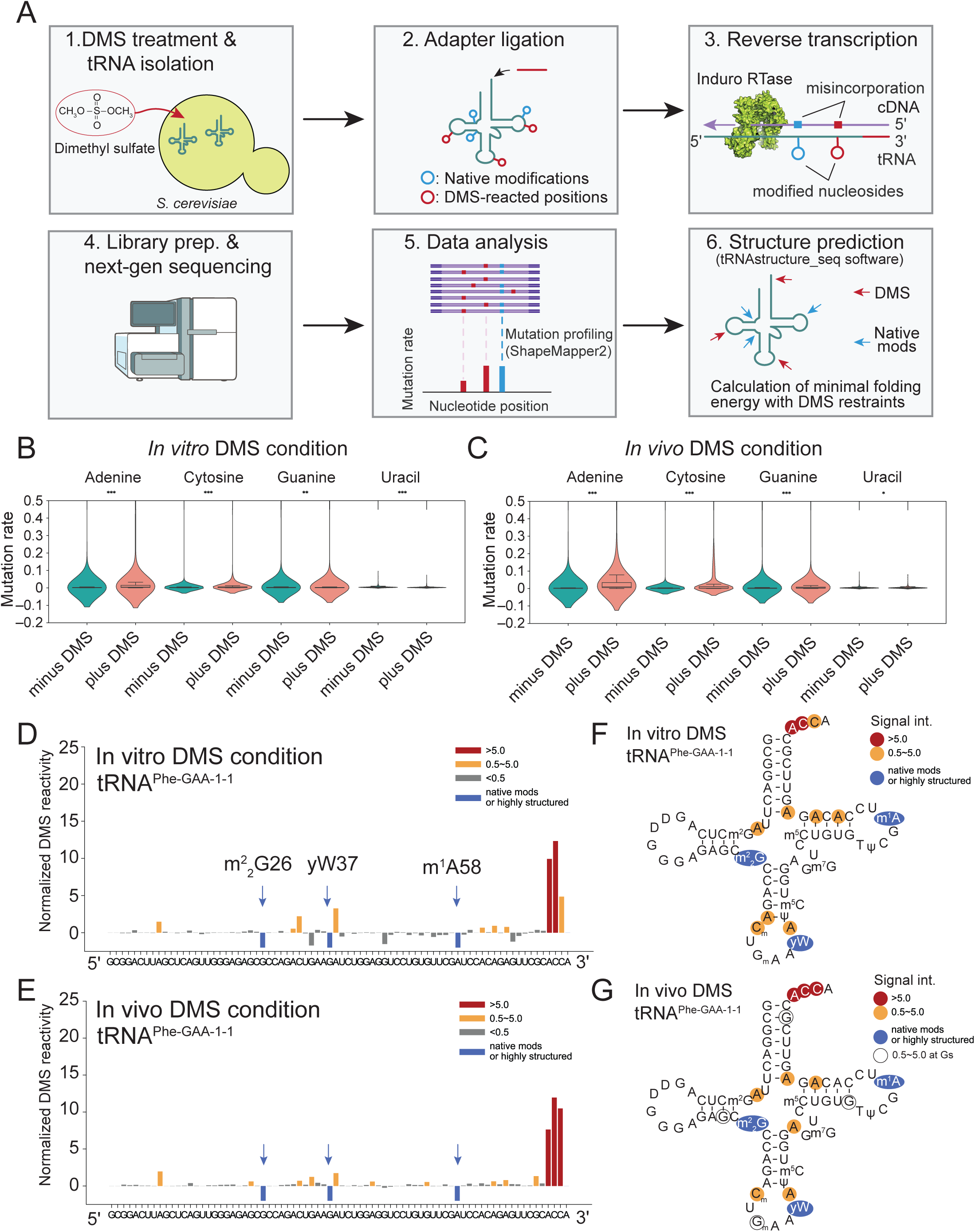
tRNA Structure–seq detects DMS–induced and native modified nucleosides in tRNA in *S. cerevisiae*. (A) Schematic illustration of tRNA Structure–seq analysis is shown. The illustration of NGS sequencer is obtained from NIH BIOART SOURCE. (B and C) Mutation rates calculated from misincorporation and indel rates at each nucleotide under (B) in vitro DMS condition and (C) in vivo DMS condition, were plotted. Minus DMS and Plus DMS were statistically compared using Student’s t–test, and asterisks indicate statistical significance (***, p–value < 0.001; **, p–value < 0.01; *, p–value < 0.05). (D and E) DMS and native modification profiling were plotted onto the primary sequence of *S. cerevisiae* tRNA^Phe–GAA–1–1^ under (D) in vitro DMS condition and (E) in vivo condition, where red indicates reactivity > 5, orange indicates 0.5 < reactivity < 5, grey indicates reactivity < 0.5. Native modification sites were identified in minus DMS samples by ShapeMapper2. To distinguish native modification signals from DMS reactivities, these sites are highlighted in blue and displayed as negative values. (F and G) DMS reactivities and native modification signals were mapped on the secondary structure of *S. cerevisiae* tRNA^Phe–GAA–1–1^ under (F) in vitro DMS condition and (G) in vivo DMS condition.

We observed that the mean values of mutation rates at adenosine and cytosine residues in the plus DMS samples were significantly higher than those in the minus DMS samples while using Induro reverse transcriptase (Fig. 1B–1C). In contrast, the mean values of mutation rates at guanosines and uridines in the plus DMS samples showed *lower* reactivities than those reactivities in the minus DMS samples (Fig. 1B–1C), although a subset of DMS–modified guanosines were still detectable at a single–nucleotide level (Fig. 1D–1E). Analysis of native tRNA modifications in tRNA^Phe–GAA–1–1^ also showed a similar trend where m^7^G46 was not detectable in both in vitro and in vivo conditions, while signals from other native modifications, presumably from m^2^_2_G26, yW37, and m^1^A58, were detected with Induro (Fig. 1D–1E), similar to with Marathon reverse transcriptase (Yamagami et al. 2022) and appear as high reactivities in minus DMS samples. DMS reactivities at adenosines and cytidines were observed throughout the tRNA, with the CCA terminus showing high reactivities under both in vitro and in vivo conditions (Fig. 1D and 1E). Mapping of DMS reactivities onto the reference secondary structure of tRNA^Phe^ revealed similar patterns of DMS reactivities between in vitro and in vivo conditions (Fig. 1F and 1G), although there were a few more protections of bases in vivo, perhaps due to interaction with aminoacyl tRNA synthetases or the ribosome.

To assess whether DMS reactivity patterns correspond to the canonical tRNA cloverleaf structure, we visualized overall DMS reactivities across all tRNA species in *S. cerevisiae* (Fig. 2A–2C for in vivo DMS condition and Supplemental Fig. 4A and 4B for in vitro DMS condition). Heatmap and box plot analyses revealed that the CCA terminus and discriminator base showed high DMS reactivities under both in vitro (Supplemental Fig. 4A and 4B) and in vivo (Fig. 2A and 2C) conditions, indicating that these 3’–terminal nucleotides are predominantly unpaired. This observation is consistent with the functional importance of the CCA terminus, which is a binding site for multiple tRNA–interacting proteins, including aminoacyl–tRNA synthetases, elongation factors (eEF1A/EF–Tu), Exportin, and tRNA modification enzymes (Zhang 2024). We also observed a hyper–DMS reactive site at A58 in the T–loop (Fig. 2A), consistent with our previous observations (Yamagami et al. 2022). A58 forms a reverse Hoogsteen base pair with U54 in the T–loop, stabilizing the interaction between the D– and T–loops (Shi and Moore 2000). This reverse Hoogsteen base pairing exposes the N1 atom of A58 to the solvent, thereby promoting DMS modification (Yamagami et al. 2022). Thus, elevated DMS reactivity at A58 likely reflects a structural signature of the canonical T-loop conformation. Also, these observations indicate that A58 remains accessible in vivo and is likely to be not extensively protected by tRNA-related proteins. Consistent with this interpretation, cryo-electron microscopy analysis of the *S. cerevisiae* RNase P–tRNA complex showed that RNase P binds the tRNA from the side opposite to A58 during cleavage of the 5’–leader sequence, leaving the N1 atom of A58 solvent-exposed (Lan et al. 2018). The configuration of the complex structure therefore supports our DMS reactivity data. Note that, because *S. cerevisiae* contains native m^1^A58 modification only in a subset of tRNA species, which can be identified in minus DMS samples (Sordyl et al. 2026), DMS reactivity at A58 from those tRNAs lacking the native modification, can be still detected. Moreover, the anticodon loop and the 3′–end of the D–loop show high DMS reactivities. To determine whether these DMS modification patterns are conserved in two organisms spanning two domains of life, we compared DMS reactivity profiles between *E. coli* and *S. cerevisiae* tRNAs (Fig. 3). Although absolute DMS reactivities varied due to differences in experimental conditions, the overall DMS modification patterns were highly similar between the two organisms. In both cases, A58, the CCA terminus, the anticodon loop, and the 3′-end of the D–loop showed elevated DMS reactivities (Fig. 3), and these conserved DMS reactivity patterns are consistent with the canonical cloverleaf secondary structure of tRNA.

**Figure 2.**
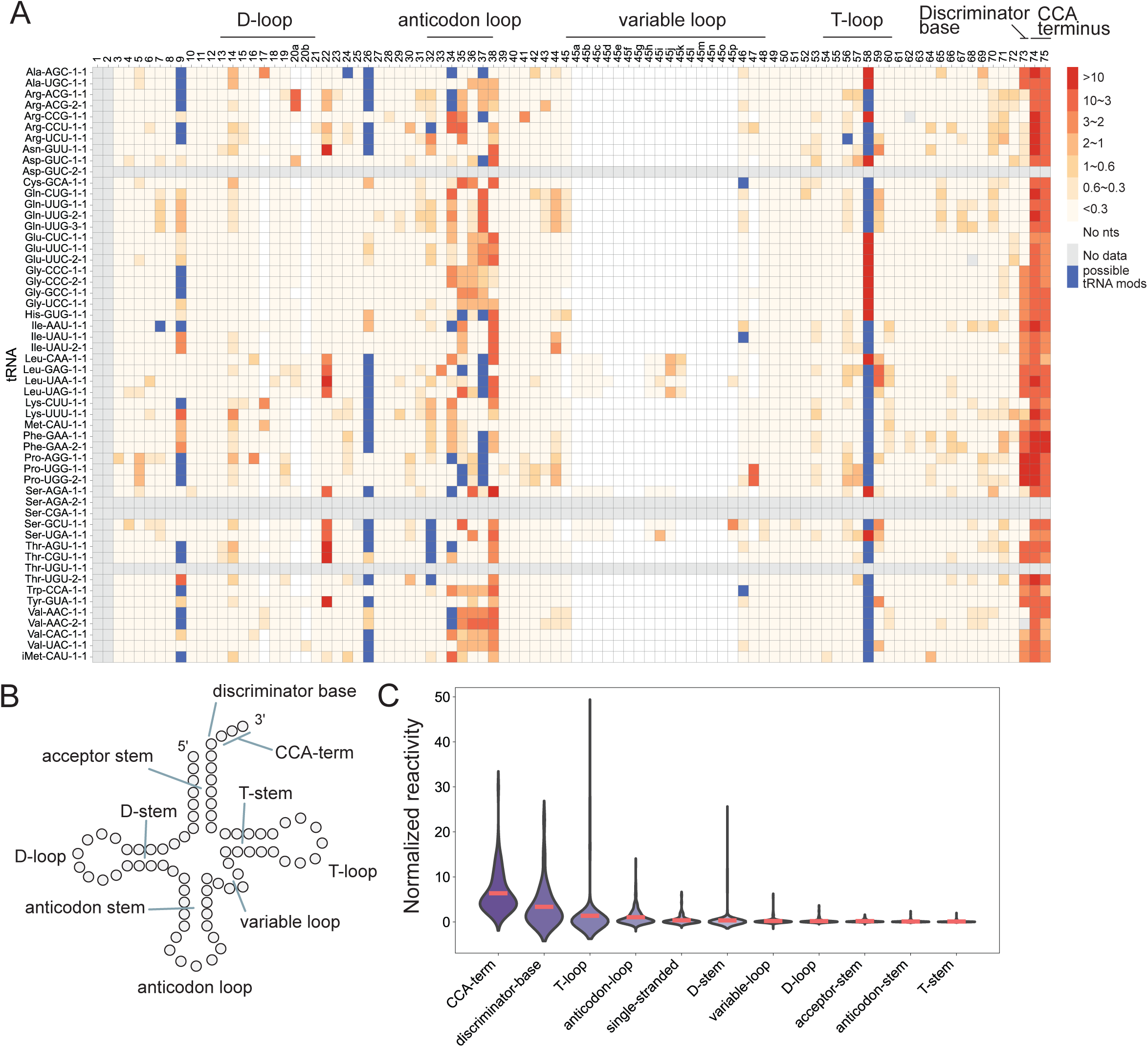
Global DMS profiling data are consistent with the canonical cloverleaf structure of tRNA in *S. cerevisiae*. (A) DMS reactivity profiles for each tRNA species were visualized. The tRNA sequences were aligned with a covariance model and then manually adjusted for the heatmap. Three successive putative native modifications identified by ShapeMapper2 were excluded from the plot, as three adjacent modifications are rarely observed in the Modomics database. The last nucleotide at position 76 is also excluded from the plot, as the nucleotide was not detected in our MaP analysis. (B) Schematic representation of the canonical cloverleaf structure of tRNA, with individual structural regions labeled. (C) The reactivities are plotted by tRNA structure region where the red line indicates the mean reactivity for each region.

**Figure 3.**
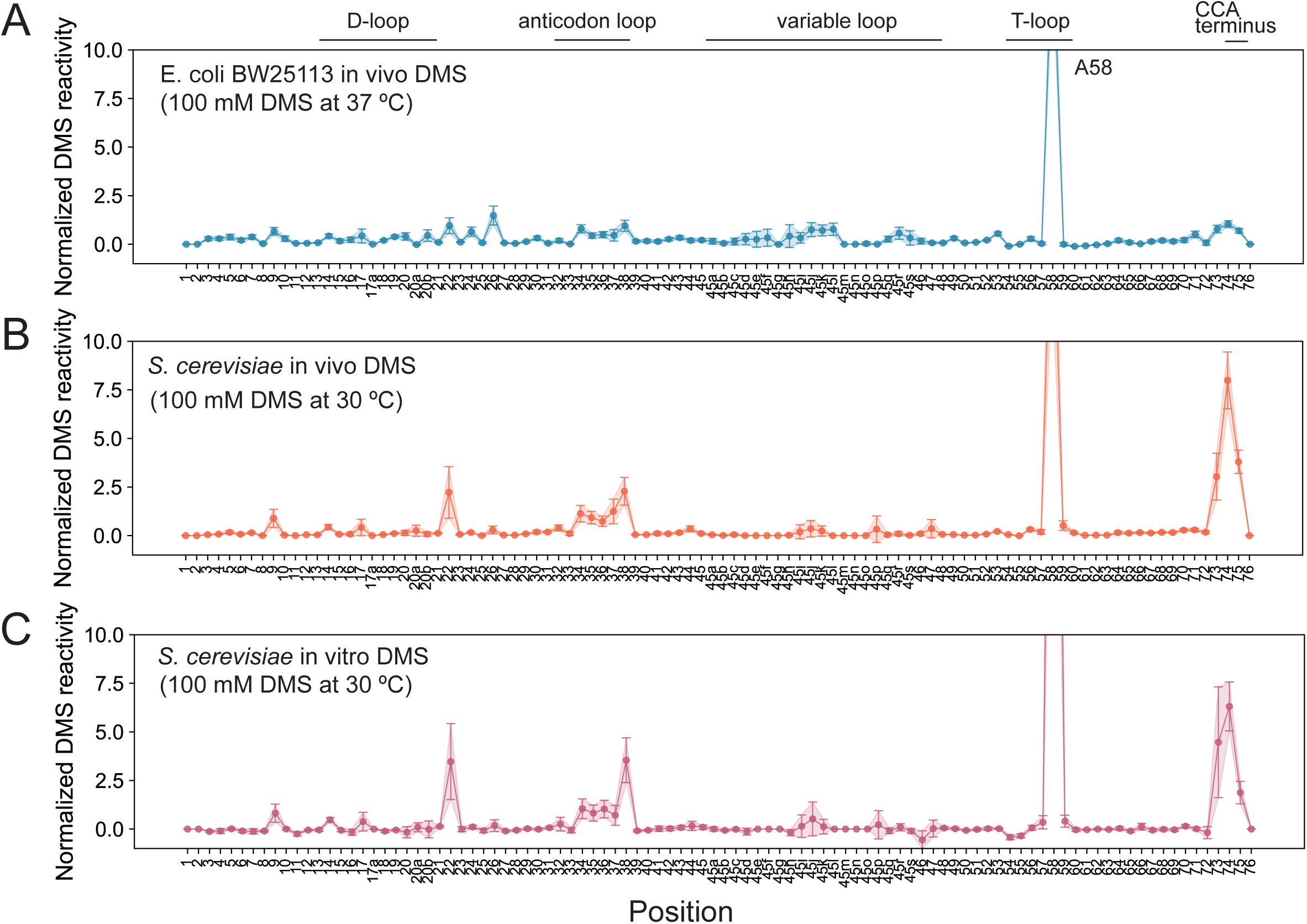
DMS modification patterns indicate that the tRNAs fold into the canonical structure in both *E. coli* and *S. cerevisiae*. (A–C) Position–specific averaged DMS reactivity profiles from three different conditions (A) *E. coli* under in vivo condition, (B) *S. cerevisiae* under in vivo DMS condition, (C) *S. cerevisiae* under in vitro DMS condition were compared. The *E. coli* dataset was obtained from source data published in our previous research (Yamagami et al. 2022) and reanalyzed in this study.

### In silico prediction of *S. cerevisiae* tRNA structure yields low prediction accuracy

Using the obtained DMS reactivity profiles, we performed secondary structure prediction of *S. cerevisiae* tRNAs (Fig. 4). As a control, we first evaluated in silico prediction without incorporating any experimental data. Reference FASTA and CT files were generated from tRNAscan–SE output to enable reproducible validation, and secondary structures were predicted using RNAstructure Fold, which performs minimum free–energy folding (Fig. 4A). The predicted structures were then compared with the reference secondary structures. Surprisingly, in silico prediction alone provided only 56.9% mean prediction accuracy (Fig. 4B), indicating that the tRNA sequences in *S. cerevisiae* are not intrinsically favored energetically to fold into the canonical cloverleaf structure using the current approach.

**Figure 4.**
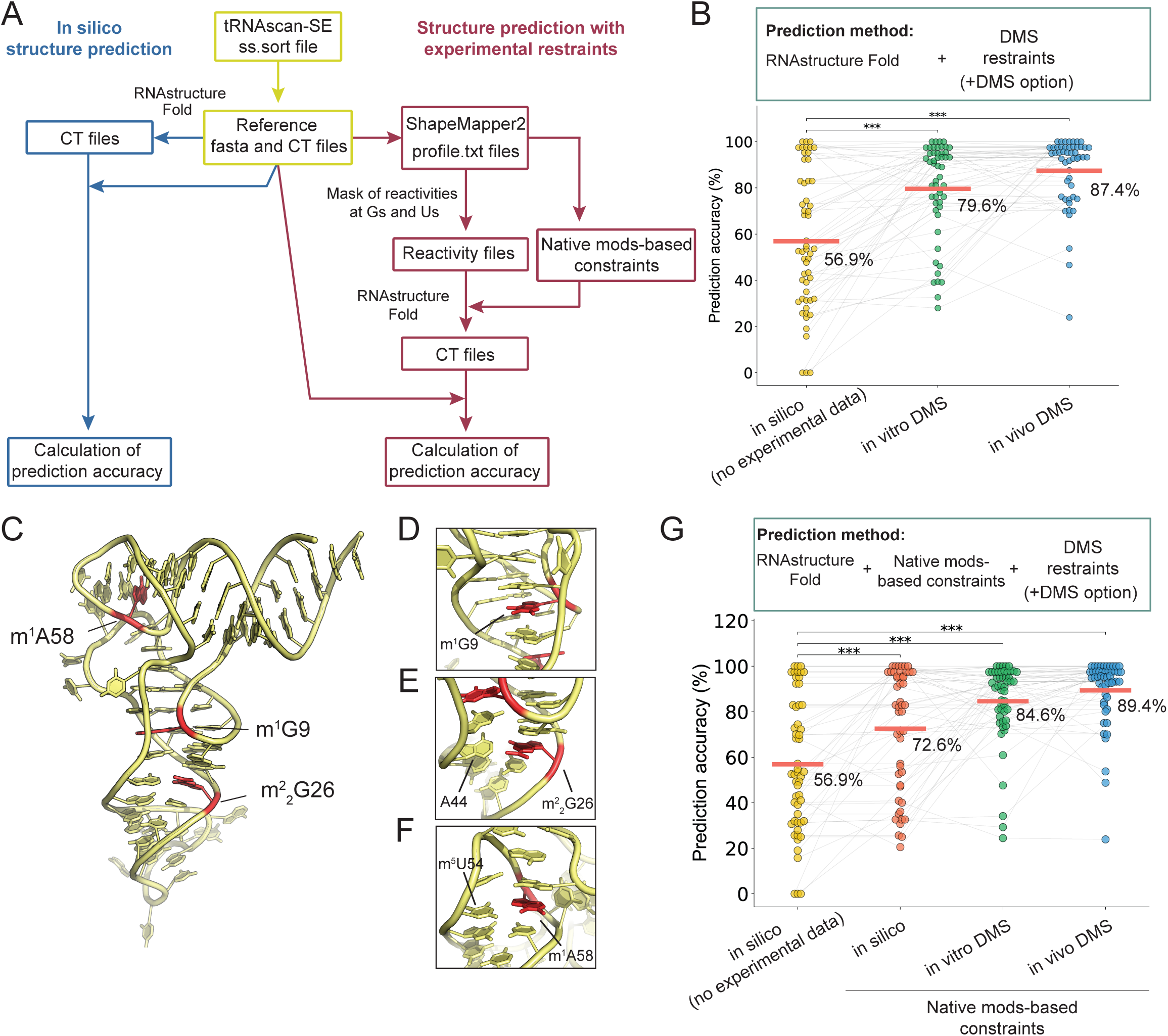
tRNA Structure–seq improves prediction accuracy of tRNA secondary structure in *S. cerevisiae*. (A) Schematic overview of tRNA Structure–seq analysis is provided. Data processing steps using tRNAscan–SE results are highlighted in yellow. In silico prediction steps are shown in blue, and in silico prediction with experimental data is shown in red. (B) Prediction accuracy of tRNA secondary structure using RNAstructure Fold with the DMS restraints option was plotted. Asterisks indicate statistical significance calculated using Student’s paired t–test (***, p–value < 0.001, in silico vs in vitro DMS, p = 1.8e–06; in silico vs in vivo DMS, p = 1.7e–08). The red line indicates the mean prediction accuracy for each condition. tRNA species lacking DMS data were excluded from the analysis. (C) Crystal structure of tRNA^Arg^ was visualized using PyMol where structure data from arginyl–tRNA synthetase–tRNA^Arg^ complex (PDB: 1F7V) was retrieved, and only the tRNA molecule was shown. The three native modifications m^1^G9, m^2^_2_G26, and m^1^A58 are highlighted in red. (D–F) Close–up views of the regions for (D) m^1^G9, (E) m^2^_2_G26, and (F) m^1^A58, are shown. (G) Prediction accuracy of tRNA secondary structure using RNAstructure Fold with both DMS restraints and native modification–based constraints, was plotted. Asterisks indicate statistical significance calculated using Student’s paired t–test (***, p–value < 0.001, in silico vs in vitro DMS, p = 1.4e–07; in silico vs in vivo DMS, p = 1.2e–09). The red line indicates the mean prediction accuracy for each condition. tRNA species lacking DMS data were excluded from the analysis.

### In silico prediction of tRNA secondary structure using DMS restraints and native modification–based constraints improves prediction accuracy

We next performed structure prediction using DMS reactivity restraints by applying the DMS option in RNAstructure Fold (Fig. 4A, left branch of red module). In this mode, DMS reactivities are incorporated as pairing–specific pseudo–energies, in which base pairing is penalized or favored according to the probability distributions of reactivity values in paired and unpaired nucleotides, respectively (Cordero et al. 2012). Incorporation of DMS restraints significantly improved mean prediction accuracy, to 79.6% under in vitro conditions and to 87.4% under in vivo conditions (Fig. 4B). However, this prediction accuracy remained insufficient for reliable tRNA secondary structure prediction. We therefore adopted an additional strategy that incorporates information from native tRNA modifications as structural constraints (Fig. 4A, right branch of red module). Specifically, we selected three conserved modifications of m^1^G9, m^2^_2_G26, and m^1^A58, which are commonly found in eukaryotic tRNAs (Sordyl et al. 2026). Because these methyl groups are installed on the Watson–Crick face, these modified nucleosides do not have an ability to form canonical Watson–Crick base pairs. Moreover, these three modification sites are located within the three–dimensional core of the tRNA molecule (Fig. 4C). Structural analysis of *S. cerevisiae* tRNA^Arg^ in complex with arginyl–tRNA synthetase revealed that m^1^G9 does not engage in base pairing (Fig. 4D) (Delagoutte et al. 2000), whereas m^2^_2_G26 forms a non–canonical base pair with A44 (Fig. 4E), and m^1^A58 forms a reverse Hoogsteen base pair with m^5^U54 in the T–loop (Fig. 4F) (Delagoutte et al. 2000; Pallan et al. 2008). These modifications were not present in all tRNAs. We thus looked for their presence in a given tRNA in our data. When signals corresponding to these native modifications were detected at G9, G26, and A58 in mutation profiling data from minus DMS samples, we utilized them as native modification–based constraints, wherein those nucleotides were prevented from forming base–pairs during structure prediction (Fig. 4G). Combinational use of DMS restraints and native modification–based constraints further improved prediction accuracy by 5 percentage points to 84.6% under in vitro conditions and by 2 percentage points to 89.4% under in vivo conditions (compare Fig. 4G to Fig. 4B). Notably, native modification–based constraints alone, i.e. with no DMS data, improved prediction accuracy by more than 12 percentage points for *S. cerevisiae* tRNAs with a p–value of 0.0003 (Supplemental Fig. 5A). To examine whether this improvement can extend beyond *S. cerevisiae*, we applied the same constraint strategy to human tRNAs using native modification information obtained from the mim–tRNAseq dataset (Behrens et al. 2021). As observed for *S. cerevisiae* tRNAs, incorporation of these constraints improved prediction accuracy by more than 8 percentage points for human tRNAs, with a p–value of 6.9e^−5^ (Supplemental Fig. 5B). Moreover, such effects can be drastic, with some tRNAs having accuracy prediction improved by 50 or even 100 percentage points just by including modification-based constraints (Supplemental Table 1). Taken together, these results suggest that conserved native tRNA modifications play a critical role in determining the canonical cloverleaf structure.

### Optimization of pseudo–free energy parameters further improves prediction accuracy of tRNA secondary structure

RNAstructure Fold incorporates chemical probing data using a pseudo–free energy calculation originally developed for SHAPE reactivities (Deigan et al. 2009). In this approach, a free energy is penalized to give a pseudo free energy when a nucleotide with high chemical probing reactivity is predicted to form base pairing, thereby filtering out those predicted structures that are inconsistent with probing data. The magnitude of this penalty is determined by Equation 1

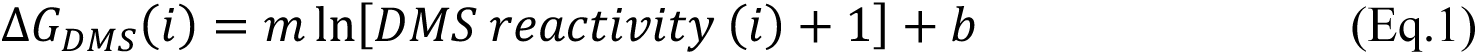

where m and b are the slope and intercept at nucleotide position i, respectively. Although this pseudo–free energy model has been widely used for RNA secondary structure prediction, the parameter values have not been optimized for tRNAs. We, therefore, tested if optimizing these pseudo–free energy parameters for tRNA structure could further improve prediction accuracy. To address this, we systematically varied the slope (m) and intercept (b) in a range of 0∼10 kcal/mol and –5∼0 kcal/mol, respectively. This analysis identified optimized parameter values of m = 2.8 kcal/mol and b = –0.8 kcal/mol for *S. cerevisiae* in vitro data (Supplemental Fig. 6A; white box), and m = 1.8 kcal/mol and b = –0.8 kcal/mol for *S. cerevisiae* in vivo data (Supplemental Fig. 6B; white box). In addition, we also revisited our previously published *E. coli* in vivo DMS data, performed the same parameter optimization, and identified optimized parameter values of m = 3.2 kcal/mol and b = –0.8 kcal/mol (Supplemental Fig. 6C; white box). We then integrated these three datasets and identified a unified parameter set of m = 2.4 kcal/mol and b = –0.8 kcal/mol (Fig. 5A; white box). The optimized parameters further improved the prediction accuracy of *S. cerevisiae* tRNA structure to 93.3% under in vivo DMS condition (Fig. 5B). In both in vitro and in vivo datasets, pseudo–free energy restraints using the unified parameter set (G_opt_) consistently outperformed the probability distribution–based model (+DMS option) by 6.7 and 3.9 percentage points, respectively. (Fig. 5C, compare Fig. 5B to Fig. 4G). Prediction accuracy for each tRNA is provided in Supplemental Table 1. For the *E. coli* in vivo dataset, the unified parameter set gave prediction accuracy comparable to that of the DMS option (probability distribution model) (Supplemental Fig. 7; green dots). We note that native modification–based constraints did not change prediction accuracy for *E. coli* tRNAs, consistent with the known absence of m^1^G9, m^2^_2_G26, and m^1^A58 modifications in this organism (Supplemental Fig. 7). Based on these results, optimized pseudo–free energy restraints (G_opt_) were used for all subsequent tRNA secondary structure predictions in *S. cerevisiae*. We provide some representative tRNA structures predicted with the optimized parameters (Fig. 5D – 5G). The tRNA Structure–seq analysis correctly predicted the canonical cloverleaf structure for tRNA^Phe–GAA^ (Fig. 5D), tRNA^Asp–GUC^ (Fig. 5E), and tRNA^Ser–UGA^ (Fig. 5F), whereas a disrupted D–loop structure was predicted for tRNA^Gln–CUG^ (Fig. 5G).

**Figure 5.**
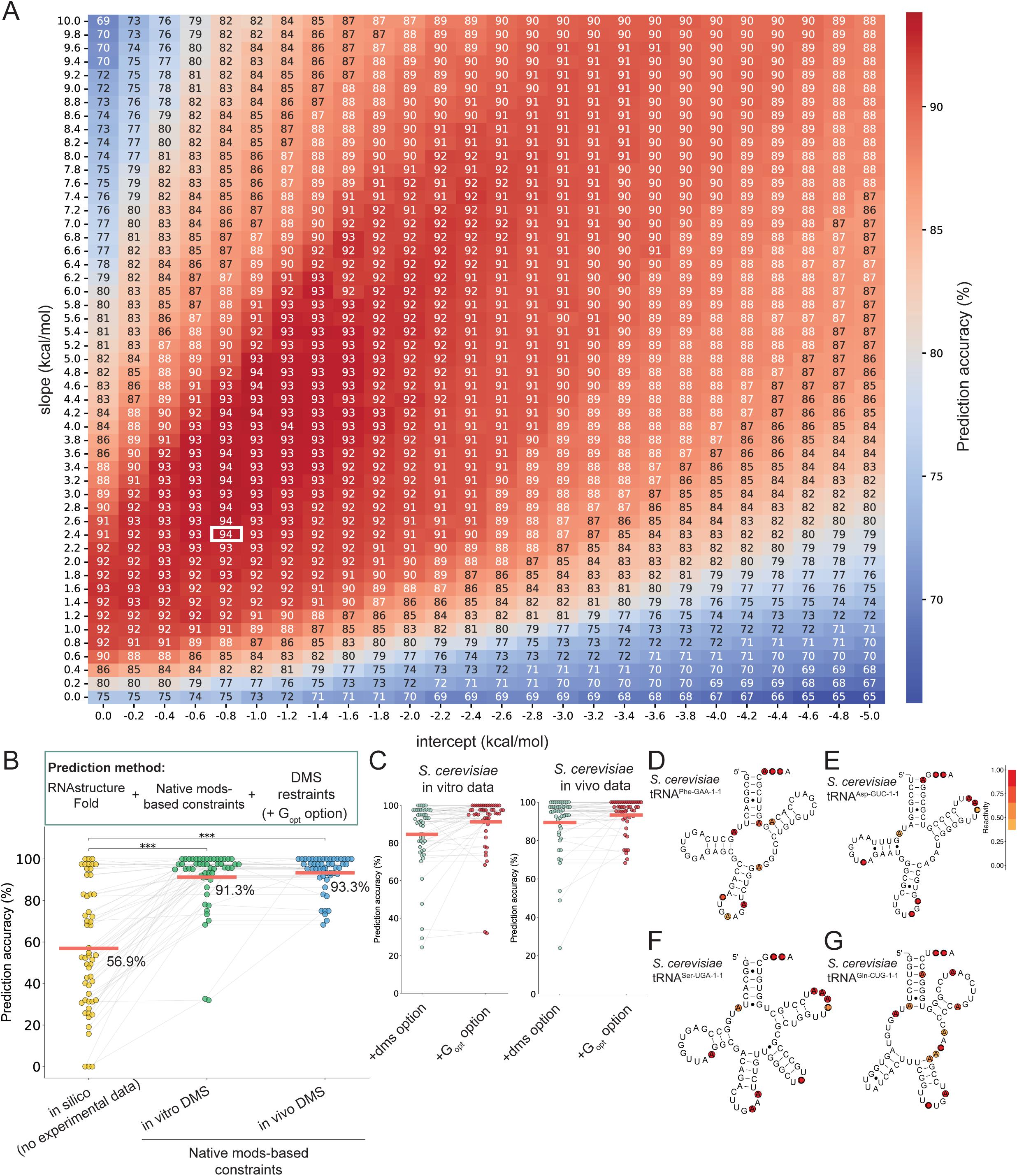
Optimization of pseudo–energy parameters further improve prediction accuracy of *S. cerevisiae* tRNA structure. (A) Pseudo–free energy parameters were systematically varied, and secondary structure prediction was performed for each parameter set. The parameters can be specified with the SHAPE option in RNAstructure Fold. Prediction accuracy was evaluated using *S. cerevisiae* in vitro and in vivo datasets, as well as *E. coli* in vivo dataset. A unified heatmap summarizing the three datasets is shown. White boxes indicate parameter sets that yielded the highest prediction accuracy. (B) Prediction accuracy of tRNA secondary structure using RNAstructure Fold with optimized DMS restraints (G_opt_ option) and native modification–based constraints were plotted. Asterisks indicate statistical significance calculated using Student’s paired t–test (***, p–value < 0.001, in silico vs in vitro DMS, p = 3.1e–11; in silico vs in vivo DMS, p = 2.6e–11). The red line indicates the mean prediction accuracy for each condition. tRNA species lacking DMS data were excluded from the analysis. (C) Comparison of prediction accuracy obtained using the DMS option and the G_opt_ option with optimized parameters for (left) in vitro DMS data and (right) in vivo DMS data. The red line indicates the mean prediction accuracy. (D–G) Representative secondary structure of *S. cerevisiae* tRNAs predicted using the optimized parameters were shown with the in vivo DMS reactivity data where (D) tRNA^Phe–GAA–1–1^, (E) tRNA^Asp–GUC–1–1^, (F) tRNA^Ser–UGA–1–1^, and (G) tRNA^Gln–CUG–1–1^ were depicted.

### Rsample analysis predicts similar global but distinct tRNA–specific conformational ensembles under in vitro and in vivo conditions

Rsample is a method that predicts multiple structures using chemical probing data (Spasic et al. 2018). Rsample generates a thermodynamic ensemble of RNA structures, estimates the expected probing reactivities from the ensemble, and iteratively reweights structure probabilities so that the predicted reactivities agree with experimental data. Using Rsample, we predicted multiple conformations of tRNA secondary structures (Fig. 6A). We first tested if the native modification–based constraints alone (i.e. no DMS data) affect the structural homogeneity of tRNA using in vivo DMS data (Supplemental Fig. 8). Interestingly, the modification–based constraints increased the fraction of tRNAs predicted to have a single conformation and decreased the fraction of tRNAs predicted to have multiple conformations (Supplemental Fig. 8; compare 8A and 8C, and 8B and 8D), indicating that the native modification–based constraints alone contribute to a structural homogeneity of tRNA.

**Figure 6.**
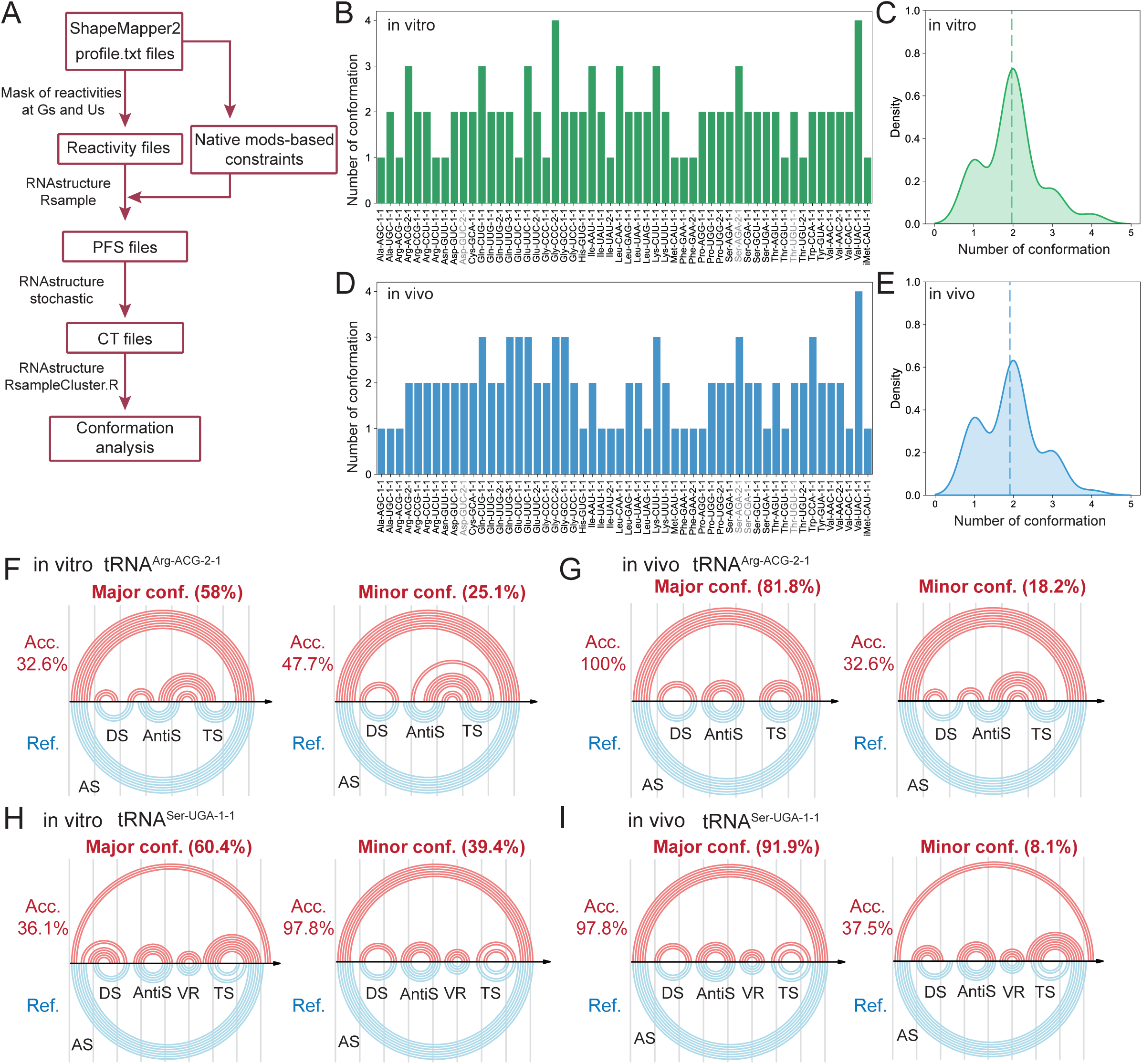
Rsample analysis shows structural homogeneity of *S. cerevisiae* tRNAs under in vitro and in vivo conditions. (A) Schematic workflow of Rsample analysis was shown. (B–E) Bar plots and histograms show the number of predicted conformations with probabilities greater than 10% for each tRNA species under (B and C) in vitro DMS conditions and (D and E) in vivo DMS conditions. The dashed line in each histogram indicates the mean number of conformations across tRNA species. (F–I) The two most probable conformations (Conf. 1 and Conf. 2) for representative tRNAs were visualized using R–chie. Numbers in parentheses and “Acc.” indicate probability of the conformation and prediction accuracy, respectively. Lines connecting nucleotides indicate predicted base pairs. Red and blue denote predicted and reference secondary structures, respectively, for (F) in vitro tRNA^Arg–ACG–2–1^, (G) in vivo tRNA^Arg–ACG–2–1^, (H) in vitro tRNA^Ser–UGA–1–1^, and (I) in vivo tRNA^Ser–UGA–1–1^. AS is acceptor stem, DS is D–stem, AntiS is anticodon stem, VR is variable region, and TS is T–stem in the reference secondary structure.

Next, we compared the tRNA conformations between in vitro and in vivo conditions, using both native modification constraints and DMS restraints. The average number of predicted tRNA conformations was similar under in vitro and in vivo DMS conditions (Fig. 6B–6E). Despite this global similarity, distinct differences were observed at the level of individual tRNAs. For example, under in vitro DMS conditions, tRNA^Arg–ACG–2–1^ populated two major conformations, neither of which corresponded to the canonical cloverleaf structure, with accuracies of just 32.6% and 47.7% (Fig. 6F). In contrast, under in vivo DMS conditions, this tRNA predominantly adopted the canonical cloverleaf structure, with a probability of 81.8% (Fig. 6G). Similarly, under in vitro conditions, tRNA^Ser–UGA–1–1^ populated two major conformations, with the major conformation predicted with an accuracy of just 36.1% and the cloverleaf–like structure predicted at a probability of only 39.4% (Fig. 6H). Under in vivo conditions, however, this tRNA folded predominantly into the cloverleaf–like structure, with a probability of 91.9% (Fig. 6I). These results indicate that, at least for these tRNAs, the canonical cloverleaf structure is more favored under in vivo conditions even though both in vitro and in vivo conditions used the same tRNAs with the same natural modifications.

### Secondary structures of *S. cerevisiae* tRNAs are robust under mild stress conditions

Finally, using the optimized tRNA Structure–seq method, we analyzed tRNA secondary structure under mild stress conditions, including heat stress, osmotic stress, and antibiotic stress (Fig. 7A). Specifically, we increased the temperature from 30 °C to 42 °C, the osmotic strength from YPD + 0% NaCl to YPD + 3% NaCl, and introduced cycloheximide (CHX) of 5 µg/mL. Growth phenotypes were monitored under each condition, and these mild stress conditions did not cause severe growth retardation (Fig. 7B – 7D). Comparison of DMS reactivity patterns between the no stress condition and the three stress conditions revealed no obvious changes in the DMS reactivity pattern, suggesting that overall tRNA secondary structures are conserved under these stress conditions (Fig. 7E). Consistent with this observation, structure prediction analyses indicated that tRNA secondary structures were largely identical across all conditions tested (Fig. 7F and 7G). Furthermore, our previous study showed a strong correlation between native tRNA modifications and mutation rates detected by MaP (Yamagami et al. 2022). We therefore compared mutation rates at G9, G26, and A58 across all conditions in the DMS minus datasets (Fig. 7H–7J). Mutation rates at G9 were significantly higher under stress conditions than under the no-stress condition, suggesting that the fraction of m¹G9-modified tRNAs increases in response to stress. In contrast, mutation rates at G26 and A58 remained largely unchanged across conditions (Fig. 7I and 7J). These changes in the modification levels were confirmed by nucleoside analysis using LC–MS/MS (Supplemental Fig. 9). These results indicate that m¹G9 levels are regulated and may be sensitive to cellular stress. Despite this overall structural robustness, a subset of tRNA species, including tRNA^Val–UAC–1–1^ and tRNA^Pro–UGG–2–1^ (highlighted by blue lines in Fig. 7F), were predicted to adopt non–canonical cloverleaf structures under specific stress conditions (Fig. 7K and 7L). Detailed comparison of individual DMS–reactive sites revealed minor changes in these tRNAs. For example, under antibiotic stress, tRNA^Pro–UGG–2–1^ gained additional DMS–reactive sites in the acceptor stem and lost a reactive site at the CCA terminus, resulting in altered local base–pairing within the acceptor stem (Fig. 7K). Similarly, tRNA^Val–UAC–1–1^ showed slight changes in DMS reactive sites within the D–loop and at the CCA terminus under antibiotic stress (Fig. 7L). Although the T–loop structure remained intact in these tRNAs, other local structural elements were perturbed under stress conditions.

**Figure 7.**
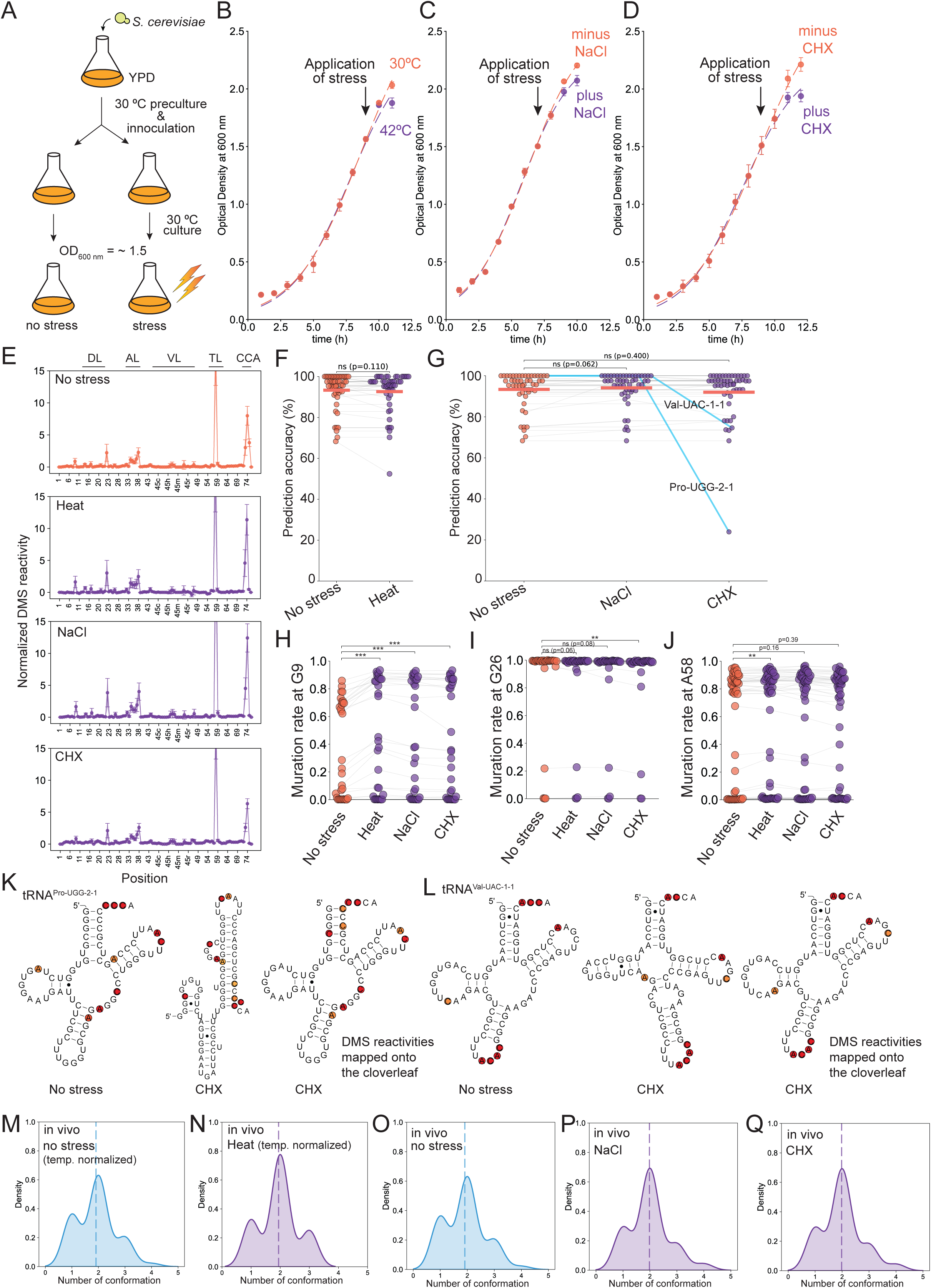
*S. cerevisiae* tRNAs are structurally stable under mild stresses. (A) Schematic illustration of the in vivo experimental design. Cells were subjected to stress at an OD_600 nm_ of ∼1.5. (B – D) Growth phenotypes under (B) heat stress, (C) osmotic stress, and (D) antibiotic stress were measured. Arrows indicate the time point at which stress was applied. For heat stress, the growth temperature was shifted to 42 °C. For osmotic stress, NaCl was added to the culture medium to a final concentration of 3% (w/v). For antibiotic stress, cycloheximide (CHX) was added to a final concentration of 5 µg/mL. The experiments were performed in three biological replicates. (E) Comparison of DMS modification patterns between the control (no stress) condition and each stress condition. Control data were replotted from Fig. 3B. The experiments were performed in three biological replicates. (F – G) Prediction accuracy of tRNA secondary structure using RNAstructure Fold with optimized DMS restraints (G_opt_ option) and native modification–based constraints under (F) osmotic and antibiotic stress conditions and (G) heat stress conditions. Statistical significance was calculated using Student’s paired t–test. The red line indicates the mean prediction accuracy for each condition. tRNA species lacking DMS data were excluded from the analysis. For heat stress conditions, DMS reactivities were temperature–normalized as described in Methods. (H-J) Mutation rates at (H) G9, (I) G26, and (J) A58, observed in the DMS minus datasets are plotted. Asterisks indicate statistical significance calculated using Student’s paired t–test (***, p–value < 0.001; **, p–value < 0.01). (K–L) Representative predicted structure of *S. cerevisiae* tRNAs with DMS reactivities under stress conditions were drawn where (K) tRNA^Pro–UGG–2–1^, (L) tRNA^Val–UAC–1–1^ were depicted. (M–Q) Histograms show the number of predicted conformations with probabilities greater than 10% for each tRNA species under (M) temperature–normalized in vivo no stress, (N) temperature–normalized in vivo heat, (O) in vivo no stress (replotted from Fig. 6E), (P) in vivo NaCl stress, (Q) in vivo cycloheximide (CHX) conditions. The dashed line in each histogram indicates the mean number of conformations across tRNA species.

Finally, Rsample analyses were performed to test whether the ensemble distributions of major tRNA conformations were affected by mild stress (Fig. 7M–7Q). The number of major conformations across all tRNA species remained unchanged, and density plots showed similar peak shapes under all conditions examined (Fig. 7M–7Q, and Supplemental Table 2). Together, these results indicate that global tRNA secondary structures in *S. cerevisiae* are highly robust and are not strongly perturbed by mild stress conditions.

## Discussion

In this study, we extended tRNA Structure–seq to the eukaryotic model organism *S. cerevisiae* and substantially improved the accuracy of tRNA secondary structure prediction by integrating optimized DMS restraints and native modification–based constraints. Our results provided a comprehensive pipeline for prediction of tRNA secondary structure in vivo and revealed that *S. cerevisiae* tRNA structures are globally stable across conditions tested.

Applying tRNA Structure–seq to *S. cerevisiae* demonstrated that this method is broadly applicable beyond bacteria and can capture conserved structural features of tRNAs in eukaryotes. DMS reactivity patterns observed in *S. cerevisiae* tRNAs were highly similar to those previously reported for *E. coli* (Yamagami et al. 2022), including high DMS reactivities at the CCA terminus, the anticodon loop, and A58 in the T–loop. These DMS reactivity patterns are consistent with the canonical cloverleaf structure of tRNA and reflect functionally important regions that are necessary to remain accessible for interactions with tRNA–related proteins such as aminoacyl–tRNA synthetases, elongation factors, and the ribosome (Rubio Gomez and Ibba 2020; Zhang 2024). Note that tRNA genes encoded in the *S. cerevisiae* genome do not contain the CCA terminus, which is added post-transcriptionally during tRNA maturation (Chan and Lowe 2016). The tRNA nucleotidyltransferase catalyzes addition of the CCA sequence at the 3’–end following removal of the 5’–leader and 3’–trailer sequences (Wolfe et al. 1996). This CCA addition is, then, required for aminoacylation in the nucleus and serves as a hallmark of nuclear export for mature tRNAs (Lund and Dahlberg 1998; Mohan et al. 1999; Grosshans et al. 2000). Since we purified full-length cDNAs during library preparation, population of the sequenced tRNA predominantly consists of mature tRNAs.

A striking finding of this study is the poor prediction accuracy of in silico *S. cerevisiae* tRNAs without any experimental constraints. Unlike *E. coli* tRNAs, which fold into canonical cloverleaf structures with relatively high accuracy of ∼80% using minimum free–energy models (Yamagami et al. 2022), *S. cerevisiae* tRNAs showed only ∼57% prediction accuracy when they are folded based on sequence and thermodynamic parameters (Fig. 4). This result strongly indicates that *S. cerevisiae* tRNA sequences are not intrinsically optimized to favor the cloverleaf structure. Thus, tRNA folding in vivo is likely to depend on additional factors, including post–transcriptional modifications, tRNA processing, and cellular context (Motorin and Helm 2010; Phizicky and Hopper 2010; Yamagami et al. 2018; Yamagami et al. 2021). These findings underscore the limitations of in silico prediction of RNA secondary structure based on thermodynamic parameters, for highly structured and extensively modified tRNAs (Lorenz et al. 2017). This is especially important in eukaryotes and archaea, which tend to have more heavily modified tRNAs (Hirata et al. 2019b; de Crecy-Lagard and Jaroch 2021; Phizicky and Hopper 2023).

A significant improvement in the prediction accuracy was achieved by incorporating native tRNA modifications as structural constraints during secondary structure prediction (Fig. 4 and Supplemental Fig. 5). Our data showed that three modifications conserved in eukaryotic tRNAs (m^1^G9, m^2^_2_G26, and m^1^A58) improved prediction accuracy when the modification information was incorporated. A similar idea was previously tested for human mitochondrial tRNA^Lys^ that possesses m^1^A9, where information on native tRNA modifications obtained from the Modomics database was used for structural constraints and improved prediction of mitochondrial tRNA structure (Helm et al. 1999; Washietl et al. 2012). This study was conducted before the establishment of high–throughput tRNA modification sequencing technologies. Recent advancements in modification–profiling techniques enable transcriptome–wide identification of native tRNA modifications (Clark et al. 2016; Behrens et al. 2021; Padhiar et al. 2024), making it possible to incorporate modification–based constraints into high–throughput tRNA structure prediction. Importantly, these modifications are located at the central core region within the L–shape structure of tRNA and are known to influence both local and global folding (Pallan et al. 2008; Lorenz et al. 2017; Hori et al. 2018; Petrosyan and Bohnsack 2024). Our results demonstrated that incorporation of modification–based constraints improves secondary structure prediction accuracy not only for *S. cerevisiae* tRNAs but also for human tRNAs (Supplemental Fig. 5), highlighting the generality of this approach and that often the effects are dramatic (Supplemental Table 1).

One possibility is that these natural modifications prevent population of non-native base pairs, thus acting as antigenic determinants. Such antigenic determinants have been shown to be important in RNA-protein interactions (Zhang and Nicholson 1997; Heinicke et al. 2009) and in recognition of tRNAs (Mohan et al. 1999; Pellegrini et al. 2012). In addition, by preventing canonical Watson–Crick base pairing, these modifications suppress alternative secondary structures competing with the canonical cloverleaf structure. The m^1^G9, m^2^_2_G26, m^1^A58 modifications encode structural information that is not specified by the primary tRNA sequence, thus, also acting as an epigenetic structural code. Of these, m^1^G9 and m^2^_2_G26 are thought to have originated in hyperthermophilic archaea such as *S. acidocaldarius*, *A. pernix*, and *T. kodakarensis*, where they contribute to the thermal stability of tRNA structure (Kuchino et al. 1982; Jackman et al. 2003; Kempenaers et al. 2010; Van Laer et al. 2016; Hori et al. 2018; Singh et al. 2018; Strassler et al. 2023). Furthermore, dimethylation also occurs at position both 26 and 27 in *A. aeolicus*, a hyperthermophilic eubacterium, and these two successive m^2^_2_G residues might have a stabilization effect on tRNA structure via hydrophobic interactions (Awai et al. 2009). While these stabilization effects on tRNA structure are not required for mesophilic organisms including *S. cerevisiae* and human, these modifications still contribute to tRNA folding. For example, disruption of these core modifications has been reported to induce structural rearrangements within the tRNA core (Steinberg and Cedergren 1995; Helm et al. 1998; Helm et al. 1999; Lorenz et al. 2017). In addition, in *S. pombe*, the double knockout strain of tRNA m^2^_2_G26 methyltransferase (Trm1) and RNA chaperon protein La showed the synthetic lethal phenotype, suggesting that this modification is required for correct folding of precursor tRNA^Ser^ (Vakiloroayaei et al. 2017). In the case of m^1^A58, the modification stabilizes T–arm structure (Hori et al. 2018). The m^1^A58 in tRNA is related to the installation of other tRNA modifications such as 5–methyl–2–thiouridine at position 54 (m^5^s^2^U54), pseudouridine at position 55 (Ψ55), and 2’–O–methylguanosine at position 18 (Gm18) in *T. thermophilus* (Tomikawa et al. 2010; Ishida et al. 2011; Yamagami et al. 2016). This tRNA modification network (or modification circuit) is also found in eukaryotes (Han and Phizicky 2018; Barraud et al. 2019; Hirata et al. 2019a; Yared et al. 2023). In addition, the m^1^A58 in tRNA^iMet^ in eukaryotes is closely related to tRNA degradation where hypomodified tRNA^iMet^ is reported to be degraded via rapid tRNA decay and the nuclear RNA surveillance systems (Kadaba et al. 2004; Kadaba et al. 2006; Tasak and Phizicky 2022). Thus, m^1^A58 modification contributes not only thermal stability of tRNA but also to quality control in eukaryotic cells (Motorin and Helm 2010). Besides these roles of m^1^G9, m^2^_2_G26, m^1^A58 modifications in tRNA stability, our Rsample analyses revealed that the modification–based constraints increase a structural homogeneity of *S. cerevisiae* tRNA (Supplemental Fig. 8). Therefore, these results support the model in which m^1^G9, m^2^_2_G26, and m^1^A58–mediated constraints act as an epigenetic structural code that suppress structural heterogeneity and bias the folding landscape toward the canonical cloverleaf structure. To fully validate this, tRNA Structure–seq analyses using modification–disrupted strain of *S. cerevisiae* are necessary.

New slopes and intercepts for pseudo-free energy calculations were identified and these improved prediction in vitro and in vivo by 6.7 and 3.9 percentage points, respectively. This suggests that tRNA may need a unique set of penalties compared to other RNAs where the original slopes and intercepts were developed. Original values had m and b of 1.8 kcal/mol and –0.6 kcal/mol (Deigan et al. 2009), whereas herein we identified a unified parameter sets of 2.4 and –0.8 for tRNA using three different datasets. The parameters for tRNA are more severe than that for other RNAs: there are bigger rewards for absence of reactivity and bigger penalties for presence of high reactivities. In other words, more weight is put on the tRNA reactivities perhaps because these RNAs fold more reliably than those in the original dataset. Further improvements can come from using G/U-specific reagents such as EDC and ETC (Mitchell et al. 2019b; Douds et al. 2024), which may help with predictions of tRNAs such as of tRNA^Gln–CUG^ (Fig. 5G), which had a disrupted D-loop, but was mostly guanosine and uridine residues and so devoid of information from DMS.

In general, we found that tRNA probed in vivo was better predicted than that probed in vitro. For instance, folding with both DMS restraints and native modification-based constraints led to mean prediction accuracy of 93.3% in vivo and 91.3% in vitro (Fig. 5B). Moreover, the more accurate prediction was often the fold with the minor population in vitro, but the fold with the major population in vivo (Fig. 5 F-I). This suggests that in vivo conditions can bias the population towards the native fold. Both preparations had the natural modifications, so this difference may be due to the denaturation of the in vitro samples at 90 °C or to the presence of proteins in vivo, although there were not significant numbers of protections found in vivo. This suggests that folding in vivo has features not captured by in vitro heating.

Overall, this study established optimized tRNA Structure–seq as a powerful and extensible platform for studying tRNA secondary structure in the three domains of life. By integrating chemical probing, native modification–based constraints, optimized folding parameters, and ensemble modeling, this method enables both high–accuracy structure prediction and detection of condition–dependent structural heterogeneity. Thus, this method is widely applicable to investigate how tRNA structure is regulated in such as developmental stages, tRNA–associated diseases, and stress environments.

## Methods

### *S. cerevisiae* strains and growth conditions

The *S. cerevisiae* BY4743 strain was used in this study. For all experiments, BY4743 cells were cultured in 100 mL of YPD medium at 30 °C. Precultured cells were inoculated into 100 mL of fresh YPD medium to an initial OD_600nm_ of ∼0.02 and grown at 30 °C with shaking. The optical density at 600 nm was monitored every hour. When OD_600nm_ reached at ∼1.8, BY4743 cells were collected by centrifugation at 8,000 xg for 15 min at 4 °C. The wet–cells were snap–frozen in liquid nitrogen and stored at –80 °C until use. For the stress experiments, the cells were subjected to stress at an OD_600 nm_ of ∼1.5. For heat stress, the growth temperature was shifted to 42 °C. For antibiotic stress, cycloheximide was added to the culture medium to a final concentration of 5 µg/mL. For osmotic stress, sodium chloride was added to the culture medium to a final concentration of 3% (w/v). The experiments were performed in biological triplicates (n= 3).

### Preparation of tRNA fraction

Total RNA was extracted from 0.1 – 0.2 g of BY4743 cells using TRIzol (ThermoFisher Scientific) according to the manufacturer’s instructions. The total RNA was separated by a 10% denaturing PAGE gel (10% polyacrylamide gel, 7 M urea, 1xTBE) at 250 V for 30 – 40 min at room temperature, and then, the RNA was visualized by SYBR gold staining. The tRNA fraction was excised from the gel, eluted in TEN250 buffer containing 10 mM Tris–HCl pH 7.6, 1 mM EDTA, and 0.001% SDS, and recovered by ethanol precipitation as described previously (Yamagami and Hori 2023b; Matsuda et al. 2024; Matsuda et al. 2025; Yamagami et al. 2025). RNA concentration was determined by measuring UV absorbance at 254 nm. The experiments were performed in biological triplicates (n= 3).

### In vitro DMS reaction

Purified tRNA was denatured at 90 °C for 3 min and allowed to cool to room temperature for 10 min. In vitro DMS modification was performed in a 50 µL reaction mixture containing 50 mM HEPES–KOH (pH 7.6), 100 mM KCl, 1 mM MgCl₂, 8 µM tRNA, and 100 mM DMS at 30 °C for 3 min. For minus DMS reactions, an equal volume of RNase–free water was added instead of DMS. The reaction was quenched by the addition of 50 µL of 2.5 M dithiothreitol (DTT). The modified RNAs were recovered by ethanol precipitation. In vitro DMS reactions were performed in biological triplicates (n = 3).

### In vivo DMS reaction

In vivo DMS modification was performed in 20 mL of BY4743 culture at an OD₆₀₀ of ∼ 1.5 at 30 °C for 3 min with shaking started by adding DMS to a final concentration of 100 mM. The reaction was quenched by the addition of 0.146 g DTT and incubated for an additional 2 min at 30 °C. For minus DMS samples, an equal volume of water was added instead of DMS. Cells were collected by centrifugation at 8,000 xg for 15 min at 4 °C, and the tRNA fraction was prepared as described above. In vivo DMS reactions were performed in biological triplicates (n = 3).

### Library preparation

Library preparation for tRNAseq was performed as described previously with minor protocol changes (Yamagami et al. 2022; Meyer et al. 2023a; Yamagami and Hori 2023b; Yamagami and Hori 2023a). First, tRNA deacylation was performed in a 100 µL reaction containing 500 mM Tris–HCl (pH 9.0) and ∼ 10 µg tRNA at 37 °C for 60 min, and tRNA was recovered by ethanol precipitation. Next, dephosphorylation was carried out in a 10 µL reaction containing 1x rCutSmart buffer (New England Biolabs), 1 U of recombinant Shrimp Alkaline Phosphatase (rSAP; New England Biolabs), and deacylated tRNA at 37 °C for 30 min. Then, rSAP was heat–inactivated at 65 °C for 5 min. The tRNA was recovered by ethanol precipitation. Adapter DNA oligonucleotides were adenylated in a 10 µL reaction containing 1x DNA adenylation buffer (New England Biolabs), 200 µM ATP, 40 µM adapter oligonucleotide, and 10 µM Mth RNA ligase (New England Biolabs) at 65 °C for 60 min. The Mth RNA ligase was heat–inactivated at 85 °C for 5 min. Adapter DNA oligonucleotides containing three different barcodes corresponding to each biological replicate, were used. The DNA sequences used for the library preparation were listed in Supplemental Table 3.

Adenylated oligonucleotides were ligated to the 3’–end of tRNAs in a 20 µL reaction containing 1x T4 RNA ligase reaction buffer (New England Biolabs), 25% PEG8000 (New England Biolabs), 6 µM adenylated oligonucleotide, ∼ 2 µM deacylated tRNA, and 200 U of T4 RNA ligase 2, truncated K277Q (New England Biolabs) at 16 °C for 16 hours. After the reaction, replicates were pooled, purified by a 10% denaturing PAGE gel, and recovered by ethanol precipitation.

Next, reverse transcription was performed using either Induro reverse transcriptase (New England Biolabs) in a 10 µL reaction containing 50 mM Tris–HCl (pH 8.3), 75 mM KCl, 2 mM MnCl_2_, 5 mM DTT, 1.25 mM dNTPs, 125 nM RT primer, <500 ng ligated tRNA, and 100 U reverse transcriptase, at 42 °C for 16 hours. After the RT reaction, 2 µL of 2 M NaOH was added to the mixture, and template RNA was degraded at 95 °C for 3 min. complementary DNA was purified by a 10% denaturing PAGE gel. The sequence of RT primer was listed in Supplemental Table 3.

Circularization of cDNA was performed in a 15 µL reaction containing 1x Circligase buffer (Lucigen), 2.5 mM MnCl_2_, 1 M Betaine, 50 µM ATP, 2.5 U CircLigase ssDNA Ligase, and cDNA, at 60 °C for 2 hours. Finally, indexing PCR was performed in a 75 µL reaction containing 1x buffer for KOD plus NEO (TOYOBO CO., LTD.), 1.5 mM MgSO_4_, 0.2 mM each dNTPs, 0.3 µM indexing forward/reverse primers, 6 µL circularized cDNA, 1.5 U KOD plus Neo (TOYOBO CO., LTD.). The PCR was performed with the following program: 98 °C for 30 sec; 12 cycles of 98 °C for 10 sec, 60 °C for 10 sec, 68 °C for 15 sec; 68 °C for 5 min; 20 °C hold. PCR products were separated by an 8% native PAGE gel in 1x TBE running buffer at 180 V for 1 h, extracted from the gel, and recovered by NucleoSpin Gel & PCR Clean–up (Macherey–Nagel). The sequences of indexing primers are listed in Supplemental Table 3. Library quality was assessed using an Agilent 2100 Bioanalyzer and ABI 7500 Real–Time PCR system. Libraries were sequenced on an Illumina NextSeq platform (20% PhiX, 150–cycle single–end sequencing, Mid Output flow cell) or an Element AVITI system (15% PhiX, 150 cycles, Cloudbreak FS Medium Output kit) using the standard Illumina sequencing primer.

### Data analysis

Reference tRNA sequence was obtained from the genomic tRNA database or predicted by tRNAscan–SE 2.0 (Chan and Lowe 2016; Chan et al. 2021). Reference tRNA sequences and their secondary structures were generated from tRNAscan–SE 2.0 output using custom Python scripts. The tRNAscan–SE 2.0 program predicts the tRNA secondary structure using the covariance model (Eddy and Durbin 1994). For those tRNA genes lacking the CCA terminus, the CCA region was appended to the 3’–end of tRNA genes.

The 3’–adapter sequence was removed from NGS reads by Cutadapt with quality trimming (–q 30) (Martin 2011). In the adapter trimming step, NGS reads from three biological replicates were demultiplexed by adapter barcode sequences (Supplemental Table 3). Then, DMS reactivity was calculated with ShapeMapper 2 (Busan and Weeks 2018) with individual normalization (––indiv–norm) and minimal mutation separation (––min–mutation–separation 1) options. Since DMS reactivity data were reproducible in the three biological replicates, we combined the fastq from three replicates into one file. Then, DMS profiling data was re–analyzed with ShapeMapper2. DMS profiling data was then used for tRNA structure prediction using a custom tRNA Structure–seq Python script incorporating RNAstructure software (Reuter and Mathews 2010). The script can be obtained at GitHub (https://github.com/Ryota-Yamagami/tRNAstructure_seq_v2). For structure prediction incorporating native tRNA modification–based constraints, information on native tRNA modifications was obtained from ShapeMapper 2 or mim–tRNAseq output (Behrens et al. 2021). In this study, m^1^G9, m^2^_2_G26, and m^1^A58 were used as structural constraints, as these modifications disrupt Watson–Crick base pairing and are located in single–stranded regions in the canonical tRNA cloverleaf structure. When signals corresponding to these native modifications were detected, the corresponding nucleotides in the reference FASTA sequences were converted to lowercase. RNAstructure is case–sensitive and regards lowercase letters as nucleotides that are constrained to be unpaired at those positions (Reuter and Mathews 2010). The revised FASTA files were used as an input sequence for structure prediction with RNAstructure Fold (Reuter and Mathews 2010). To optimize pseudo–free energy calculations, the slope (m) and intercept (b) parameters in Equation (1) were systematically varied. Predicted tRNA secondary structures were visualized using R2easyR and R2R software, or R–chie (Weinberg and Breaker 2011; Tsybulskyi et al. 2020; Sieg et al. 2021). Rsample was used for predicting multiple tRNA conformations from DMS profiling data (Spasic et al. 2018) where we analyzed the multiple conformations with probability greater than 10%. Temperature normalization of DMS reactivity was performed as described previously (Yamagami et al. 2022). In brief, ShapeMapper normalizes the DMS reactivity at each nucleotide position divided by the average DMS reactivity of the top 10% reactive nucleotides in a tRNA species. The temperature–normalized reactivity was calculated from the normalized DMS reactivity from ShapeMapper2 using our previous normalization method (Su et al. 2018; Yamagami et al. 2022).

### Nucleoside analysis by mass spectrometry

Twenty picomole of tRNA was digested into a nucleotide 5’-monophosphate (pN) with 0.03 units of nuclease P1 (Fujifilm Wako Pure Chemical) in 10 mM NH_4_OAc (pH 5.3) at 37 °C for 60 min, followed by adding 0.04 units of phosphodiesterase I (Worthington Biochemical) in 50 mM ammonium bicarbonate and incubating at 37 °C for 60 min. The resulted nucleotides were then dephosphorylated by adding 0.03 units of bacterial alkaline phosphatase (*E. coli* C75, Nippon Gene) and incubating at 37°C for 60 min. The nucleosides were separated using a Hypersil GOLD aQ C18 column (150 × 2.1 mm, 1.9 µm, ThermoFisher Scientific) equipped with a guard cartridge (ThermoFisher Scientific) and analyzed using an UltiMate 3000 HPLC system coupled to a Q Exactive Orbitrap mass spectrometer (ThermoFisher Scientific). The solvent system comprised 5 mM NH*4*OAc (pH 5.3) and acetonitrile (ACN) with a multi-step gradient (1–11% ACN from 0 to 6 min, 11–24% ACN from 6 to 12 min, 24–90% ACN from 12 to 15 min, and 90% ACN from 15 to 25 min) at 100 µl/min. The column was then re–equilibrated at 1% ACN from 25 to 34 min at 300 µL/min and 1% ACN from 34 to 35 min at 100 µL/min. Positive ion scanning ranged from 105 to 700 *m/z*. For relative quantification, peak areas of modified bases were obtained from extracted-ion chromatograms and normalized to the peak area of cytosine using Qual Browser bundled in Xcalibur (ThermoFisher Scientific). The experiments were performed in biological triplicates (n= 3).

## Supplemental Material

Supplemental materials are available for this article.

## Acknowledgement

The authors thank Dr. Naohito Tokunaga at the Advanced Research Support Center (ADRES at Ehime University) and Dr. Kohei Nishino at Tokushima University for assistance with the next–generation sequencing experiments and LC–MS experiments, respectively. We are grateful to the lab member. This work was supported by the JAPAN Society for the Promotion of Science (JSPS) (22K15035 and 25K18406 for R.Y.), Takeda Science Foundation (2023038199 for R.Y.), Terumo Life Science Foundation (2023tkh4 for R.Y.), the Naito Foundation (2024_206 for R.Y.), the Uehara Memorial Foundation (202410178 for R.Y.), the Sasakawa Scientific Research Grant from The Japan Science Society (2025-4046 for T.M.), and the National Institutes of Health (NIH) grant (R35GM127064 for P.C.B.).

## Data Availability

All sequencing data obtained in this study are available at the DNA Data Bank of Japan (DDBJ) Sequence Read Archive (SRA), under the accession numbers DRR902553-DRR902564. The *E. coli* DMS data has been available at https://scholarsphere.psu.edu/resources/5f58a17e-074e-4518-af5f-bb86686b1a68. All data used in this study are available from the corresponding author upon request.

## Competing interests

The authors declare no competing interests.

## Notes

### Competing Interest Statement

The authors have declared no competing interest.

### Summary of Updates

We have performed additional experiments and revised the manuscript accordingly. Supplemental Fig. 9 has been added, and the author list and affiliations have been updated.

