## Supplemental Figures for "Optimized tRNA structure–seq reveals robust tRNA secondary structures in *S. cerevisiae* under mild stress conditions"

Komei Yanagihara^1^, Futa Konishi^1^, Teppei Matsuda^1^, Akira Hirata^2^, Hiroyuki Hori^1^, Philip C. Bevilacqua^3,4,5^, and Ryota Yamagami^1,*^

^1^Department of Applied Chemistry, Graduate School of Science and Engineering, Ehime University, 3 Bunkyo-cho, Matsuyama, Ehime 790-8577, Japan

^2^Department of Natural Science, Division of Science and Technology, Graduate School of Sciences and Technology for Innovation, Tokushima University, 2-1 Minamijosanjima-cho, Tokushima, Tokushima 770-8506, Japan

^3^Department of Chemistry, Pennsylvania State University, University Park, PA 16802, USA

^4^Center for RNA Molecular Biology, Pennsylvania State University, University Park, PA 16802, USA

^5^Department of Biochemistry and Molecular Biology, Pennsylvania State University, University Park, PA 16802, USA

*** Corresponding author:**

Ryota Yamagami

Address: 3 Bunkyo-cho, Matsuyama, Ehime 790-8577, Japan

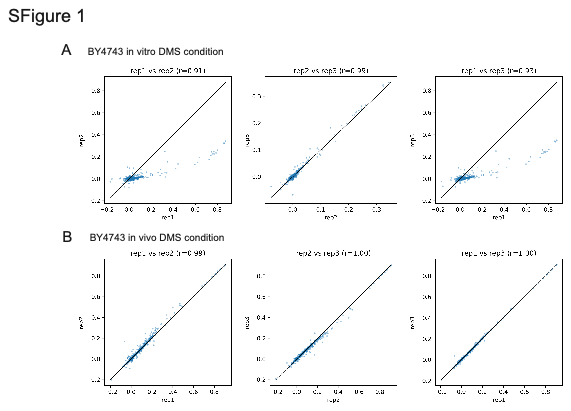

**Supplemental Fig. 1 tRNA Structure–seq datasets used in this study showed high reproducibility.** (A – B) Normalized reactivities were plotted to assess correlations between biological replicates for (A) *S. cerevisiae* BY4743 in vitro DMS conditions and (B) BY4743 in vivo DMS condition. Pearson correlation coefficient (R) values were determined by linear regression.

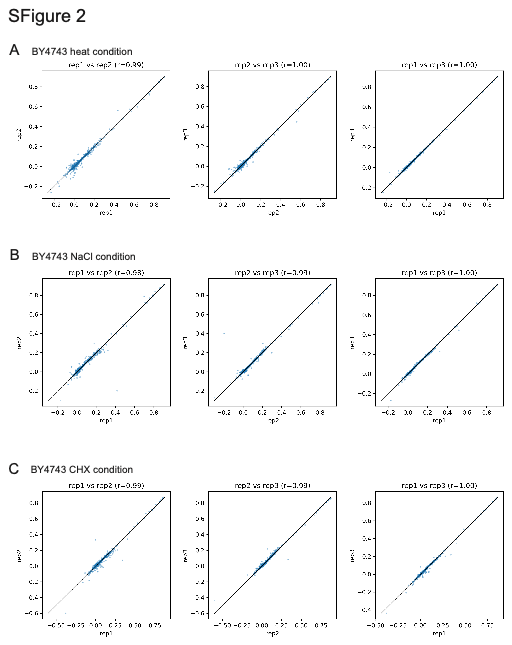

**Supplemental Fig. 2 tRNA Structure–seq datasets obtained in the mild stress conditions showed high reproducibility.** (A – C) Normalized reactivities were plotted to assess correlations between biological replicates for (A) *S. cerevisiae* BY4743 heat condition, (B) BY4743 NaCl condition, and (C) BY4743 antibiotic condition where CHX means cycloheximide. Pearson correlation coefficient (R) values were determined by linear regression.

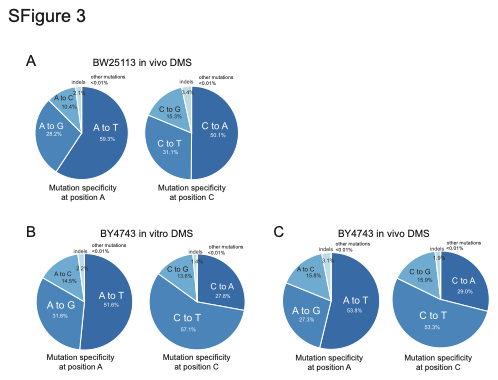

**Supplemental Fig. 3 Marathon and Induro reverse transcriptases showed similar mutation specificities.** (A – C) Mutation specificities at positions A and C were calculated from mutation profiling data generated by ShapeMapper2 for (A) *E. coli* BW25113 in vivo DMS condition, (B) yeast BY4743 in vitro DMS condition, and (C) BY4743 in vivo DMS condition. The *E. coli* dataset was obtained from source data published in our previous research (Yamagami et al. 2022) and reanalyzed in this study.

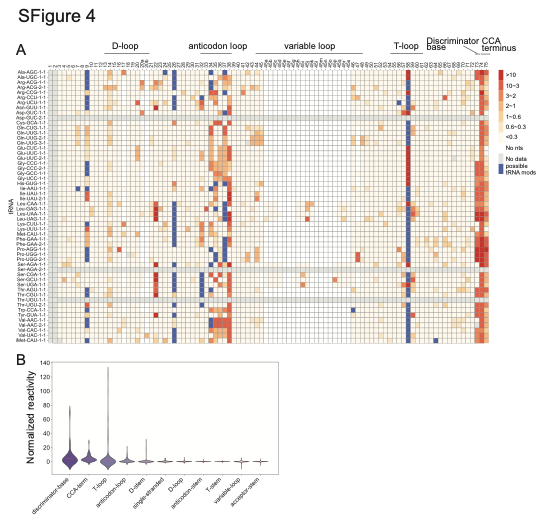

**Supplemental Fig. 4 Global DMS profiling data from in vitro DMS condition are consistent with the canonical cloverleaf structure of tRNA.** (A) DMS reactivity profiles obtained from in vitro DMS condition were visualized for each tRNA species. Three successive putative native modifications identified by ShapeMapper2 were excluded from the plot, as three adjacent modifications are rarely observed in the Modomics database. The last nucleotide at position 76 is also excluded from the plot, as the nucleotide was not detectable in our MaP analysis. (B) The reactivities are plotted by tRNA structure region where the red line indicates the mean reactivity for each region.

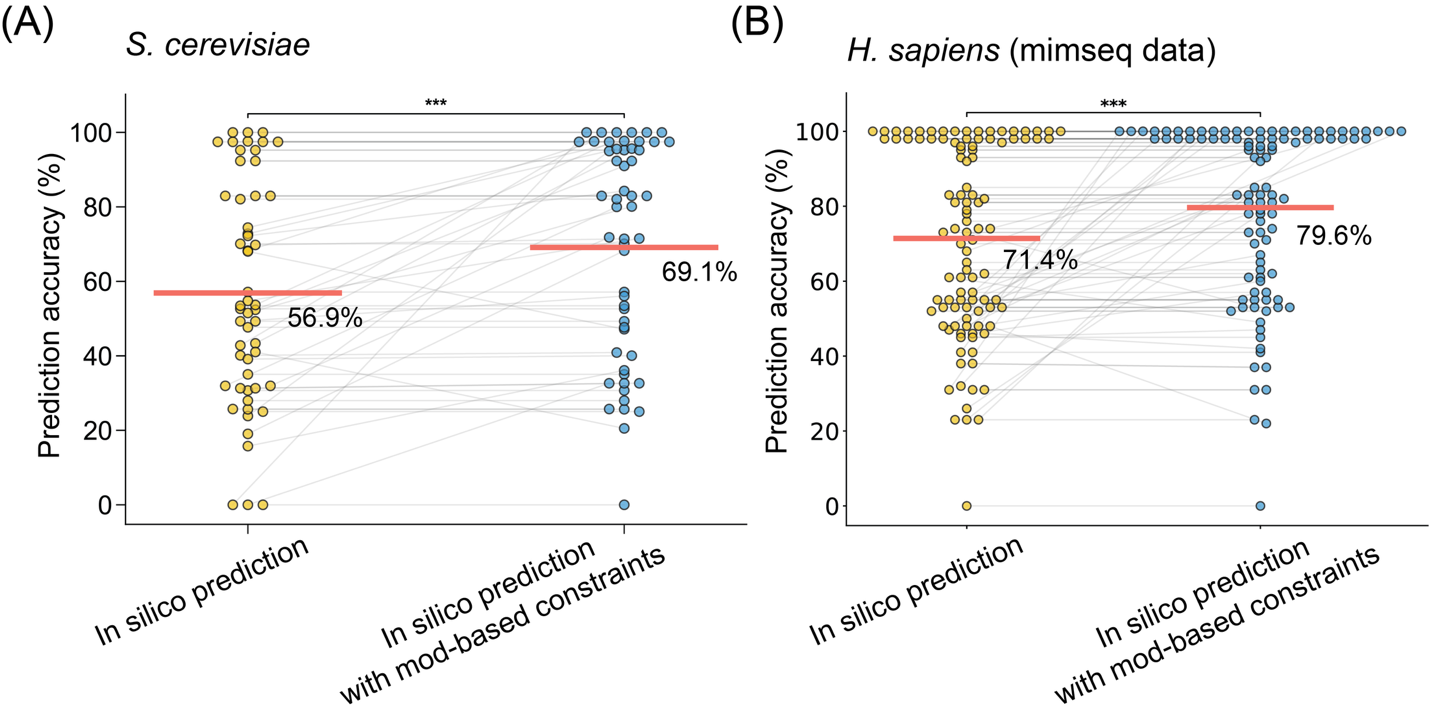

**Supplemental Fig. 5 Native modification-based constraints significantly improve prediction accuracy of tRNA secondary structure.** (A) Structure prediction was performed with/without the native modification–based constraints for *S. cerevisiae* using in vivo DMS data. The red line indicates the mean value for prediction accuracy. Asterisks indicate statistical significance calculated using Student’s paired t–test (***, p–value < 0.0003). (B) Structure prediction was performed with/without the native modification–based constraints for *H. sapience* using published mimseq data (Behrens et al. 2021). The red line indicates the mean value for prediction accuracy. Asterisks indicate statistical significance calculated using Student’s paired t–test (***, p–value < 6.9e^-5^).

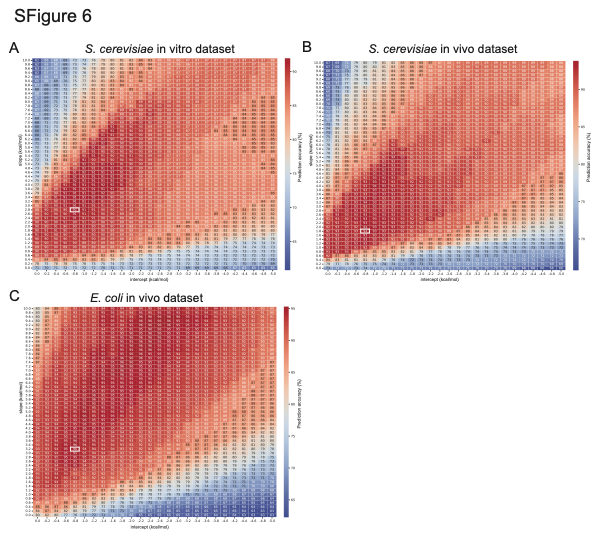

**Supplemental Fig. 6 Optimization of pseudo–energy parameters was performed for *S. cerevisiae* and E. coli tRNAs.** (A–C) Pseudo–free energy parameters were systematically varied, and secondary structure prediction was performed for each parameter set. The parameters can be specified with the SHAPE option in RNAstructure Fold. Prediction accuracies for (A) *S. cerevisiae* in vitro DMS condition, (B) *S. cerevisiae* in vivo DMS condition, and (C) *E. coli* in vivo DMS condition, are shown as a heatmap. White boxes indicate parameter sets that yielded the highest prediction accuracy. The *E. coli* dataset was obtained from our previous report (Yamagami et al. 2022).

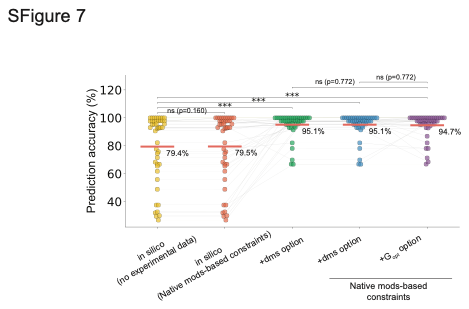

**Supplemental Fig. 7 Comparison of secondary structure prediction of tRNAs using in vivo DMS dataset from *E. coli* BW25113 strain.** Prediction accuracy was compared across different conditions: in silico prediction without experimental data, prediction with native modification–based constraints, prediction using the dms option, prediction using the dms option combined with native modification–based constraints, and prediction using the G_opt_ option with optimized parameters and native modification–based constraints. The red line indicates the mean prediction accuracy for each condition. tRNA species lacking DMS data were excluded from the analysis.

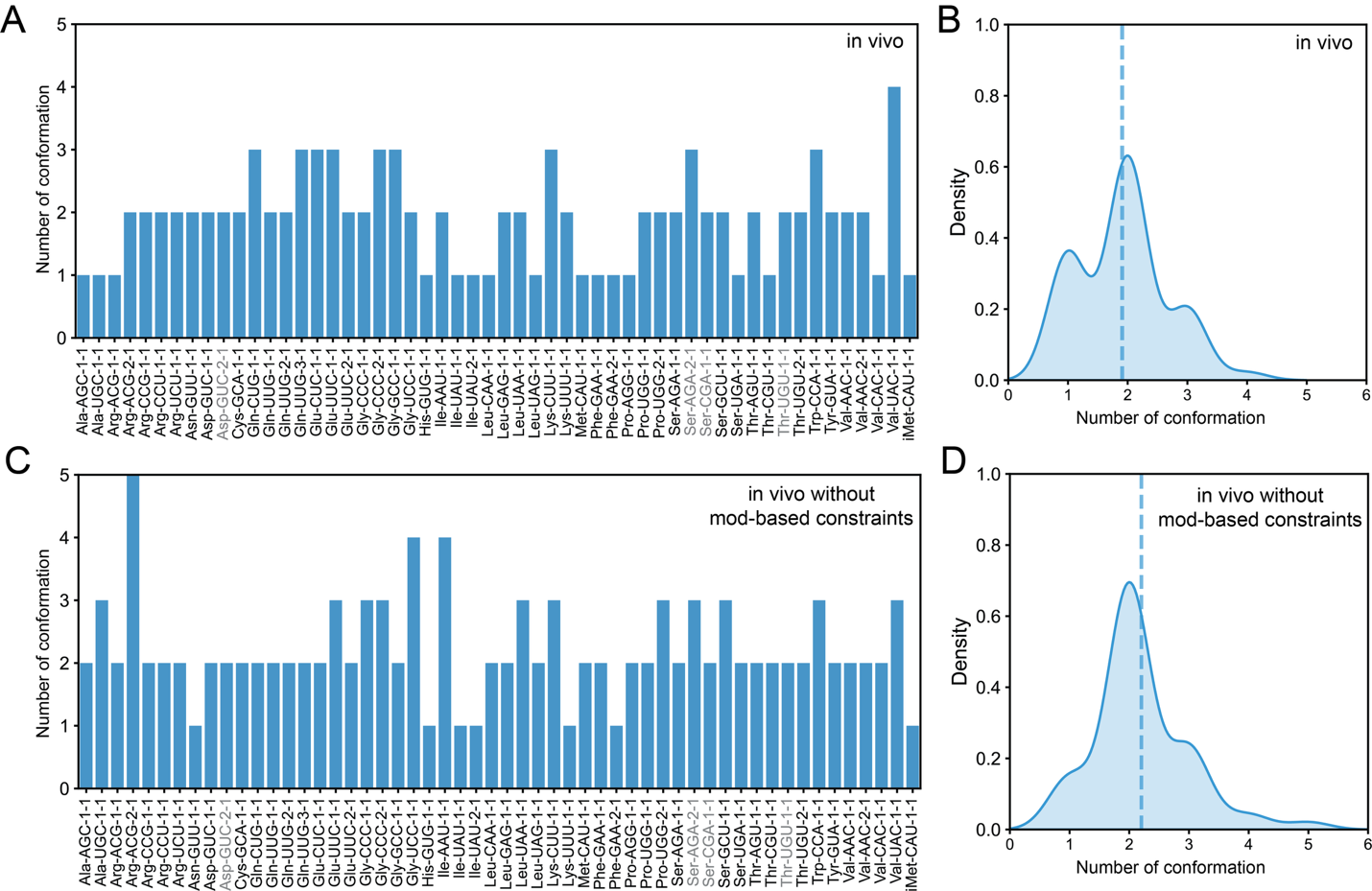

**Supplemental Fig. 8 native modification–based constraints increase structural homogeneity of yeast tRNAs.** (A–D) Bar plots and histograms show the number of predicted conformations with probabilities greater than 10% for each tRNA species under (A, B) in vivo DMS conditions and (C, D) in vivo DMS conditions combined with native modification–based constraints. The dashed line in each histogram indicates the mean number of conformations across tRNA species.

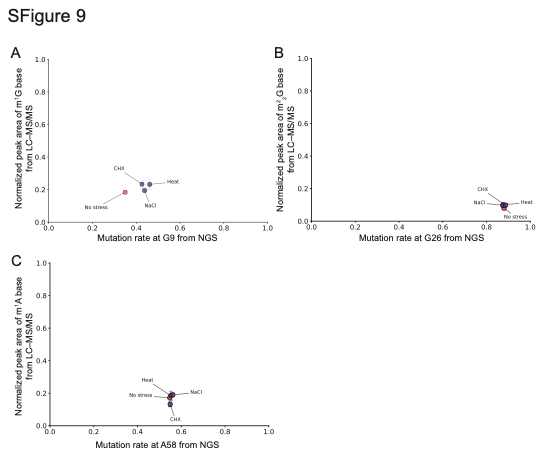

**Supplemental Fig. 9 m^1^G modification levels increase under mild stress conditions.** (A–C) Relative quantification by mutation profiling was cross-validated by LC–MS/MS experiments. Peak areas of (A) m^1^G, (B) m^2^_2_G, and (C) m^1^A bases were quantified and normalized to the peak area of cytosine. The averaged mutation rates at G9, G26, and A58 obtained from next–generation sequencing experiments were compared with normalized peak areas of m^1^G, m^2^_2_G, and m^1^A nucleosides measured by LC–MS/MS under mild stress conditions. Error bars indicate the standard deviation from three biological replicates (n = 3).
